# Single-cell phylogenomics identifies major groups of marine eugregarine parasites (Apicomplexa)

**DOI:** 10.1101/2025.04.08.647885

**Authors:** Eunji Park, Ina Na, Alana Closs, Kyle Hall, Tyrel Froese, Danja Currie-Olsen, Ondine Pontier, Niels Van Steenkiste, Patrick J. Keeling, Brian S. Leander

## Abstract

Gregarines are a large group of apicomplexan parasites that infect a wide range of invertebrate hosts, including diverse and speciose groups, such as annelids and arthropods. Marine eugregarines represent the majority of gregarine diversity, but remain poorly understood, especially their deepest phylogenetic relationships. To expand knowledge of marine eugregarine diversity and their evolutionary history, we surveyed marine invertebrates, with a particular focus on annelids, across multiple locations in British Columbia, Canada. From this effort, we obtained high-quality, single-cell transcriptomes from 20 different species of marine eugregarines, including nine previously described species and 11 novel ones, which more than doubles the amount of phylogenomic data for the group. These data, which comprehensively represent the known diversity of marine gregarines in annelid hosts, allowed us to construct an expanded phylogenetic tree based on small subunit ribosomal DNA sequences and a phylogenomic tree inferred from 142 proteins and 44,802 amino acid sequences. Our analyses identified five “superfamily-level” groups of marine eugregarines infecting annelid hosts: The Ancoroidea, Lecudinoidea, Loxomorphoidea n. superfam., Paralecudinoidea n. superfam., and Belladinoidea n. superfam., with the latter three newly established in this study. These findings contribute to ongoing efforts to build a robust molecular phylogenetic framework for gregarine diversity and refine gregarine classification, supporting the recognition of 11 eugregarine superfamilies. However, some of the deepest evolutionary relationships among these superfamilies remain unresolved, highlighting the need for expanded taxon sampling to better capture the true diversity of eugregarine parasites.

## 1. INTRODUCTION

Apicomplexans are a diverse group of single-celled parasites, including several notorious pathogens such as *Plasmodium* (Hematozoa) and *Toxoplasma* (Coccidia), which cause malaria and toxoplasmosis in humans and other vertebrates (Current and Garcia, 1991; Weiss and Dubey, 2009). Due to their significant impact on human health and the economy, these parasites have been extensively studied. In contrast, gregarine apicomplexans do not infect humans but play an important ecological role due to their diversity and abundance in species-rich invertebrate hosts, such as annelids and arthropods (Desportes and Schrével, 2013; Leander, 2008; Levine, 2018). Unlike many other apicomplexans that require both intermediate and definitive hosts, gregarines have a relatively streamlined life cycle that involves a single host (Perkins et al., 2000). Infection begins when a host ingests oocysts from its environment. Inside the host, infective sporozoites emerge from the oocysts and invade the gut lining, developing into larger feeding cells, called trophozoites, that inhabit extracellular spaces, such as the intestinal lumen. As they mature, two trophozoites pair up and undergo a process called syzygy, forming a gametocyst. Within this structure, haploid gamonts mitotically divide to produce male and female gametes, which fuse to form zygotes. These zygotes develop into oocysts, undergoing meiosis to generate multiple (e.g., 4 or 8) sporozoites. Each gametocyst can contain hundreds of oocysts, which are eventually released into the environment through host defecation or post-mortem decay, continuing the infection cycle (Desportes and Schrével, 2013; Schrével, 1969).

Among the developmental stages of gregarines, trophozoites are the most conspicuous and readily encountered due to their large size, active movement and presence within the intestinal lumen of a host. As a result, species descriptions have traditionally been based on trophozoite morphology. However, classifying gregarines has been challenging due to a limited number of distinguishing morphological traits in trophozoites, morphological variation within populations of the same species, and widely scattered taxonomic literature (Clopton, 2009; Levine, 1977a; Perkins et al., 2000; Simdyanov et al., 2017). Historically, gregarines have been categorized into three major groups—Archigregarines, Eugregarines, and Neogregarines—based on life cycle and trophozoite morphology (Grassé, 1953; Perkins et al., 2000). Eugregarines were further divided into septate and aseptate forms based on the presence or absence of a transverse septum in the trophozoites (Chakravarty, 1960; Levine, 1971). However, molecular phylogenetic analyses using small subunit ribosomal RNA gene (syn. SSU rDNA) sequences have revealed that these classifications do not reflect evolutionary relationships. For instance, neogregarines were found to be polyphyletic with different lineages nested throughout eugregarines (Simdyanov et al., 2017). Similarly, septate eugregarines do not form a distinct clade and are interspersed among lineages of aseptate lineages (Simdyanov et al., 2017). While phylogenies inferred from only SSU rDNA sequences could not confirm the monophyly of gregarines and eugregarines, more recent phylogenomic studies have provided strong evidence that gregarines are monophyletic and comprise two major subclades: one consisting of archigregarines and blastogregarines, and one consisting of eugregarines (Janouskovec et al., 2019; Lax et al., 2024; Mathur et al., 2019).

Eugregarines infect a wide range of invertebrate hosts across marine and terrestrial environments (Desportes and Schrével, 2013). While SSU rDNA sequences have not been particularly effective in resolving deeper relationships between major gregarine groups, several distinct clades have been identified, leading to multiple superfamily-level taxa (Clopton, 2009; Simdyanov et al., 2017): The Actinocephaloidea Léger, 1892, Gregarinoidea Labbé, 1899, Cephaloidophoroidea Rueckert et al. 2011, Lecudinoidea Simdyanov and Diakin 2013, and Ancoroidea Simdyanov 2013. Gregarines that infect terrestrial arthropods have received additional taxonomic attention, leading to the recognition of five superfamilies: The Actinocephaloidea Léger, 1892, Stylocephaloidea Clopton, 2009, Dactylophoroidea Miroliubova et al., 2025, Stenophoroidea Clopton, 2009, and Gregarinoidea Labbé, 1899 (Miroliubova et al., 2025). However, despite their well-documented molecular and morphological diversity, only two superfamilies—the Lecudinoidea Simdyanov, 2013 and Ancoroidea Simdyanov et al., 2017 —have been established for marine eugregarines that infect annelids and other invertebrates.

Historically, the family Lecudinidae (Lecudinoidea) served as a broad taxonomic category for most marine aseptate eugregarines, but molecular phylogenetic studies have demonstrated that *Lecudina*-like gregarines are highly divergent and form multiple independent lineages (Levine, 1977a, 1976; Park and Leander, 2024a). The Lecudinoidea currently includes ten confirmed genera: *Lecudina*, *Lankesteria*, *Difficilina*, *Lithocystis*, *Pterospora*, *Urospora*, *Veloxidium*, *Undularius*, *Amplectina*, and *Sphintocystis* (Leander et al., 2006; Odle et al., 2024; Park and Leander, 2024a; Rueckert et al., 2015, 2011, 2010; Rueckert and Leander, 2008; Simdyanov, 2004; Wakeman and Leander, 2012). Ancoroidea consist of *Ancora*, *Polyrhabdina*, *Poliplicarium* and *Trollidium* (Janouskovec et al., 2019; Simdyanov et al., 2017; Wakeman, 2020). Some other lineages remain unclassified within these superfamilies and form long branches in phylogenies inferred from SSU rDNA sequences. For example, *Paralecudina polymorpha* was initially described as *Lecudina* due to morphological similarities of the trophozoites, but was later reclassified based on its distinct molecular phylogenetic position (Iritani et al., 2018b; Leander et al., 2003; Rueckert and Leander, 2010). Similarly, *Loxomorpha* and *Cuspisella*, which infect polynoid annelids, and *Trichotokara* species, found in onuphid and eunicid annelids, exhibit long-branches in phylogenies inferred from SSU rDNA sequences (Iritani et al., 2018a; Rueckert et al., 2013; Rueckert and Leander, 2010).

In this study, we investigated marine annelid hosts, along with other invertebrates such as nemerteans, echinoderms and tunicates, over a two-year period in British Columbia, Canada, to survey the diversity of marine eugregarines. We obtained 20 high quality, single-cell transcriptomes from both described and undescribed species of marine euegregarines, more than doubling the amount of phylogenomic data for the group, providing a powerful signal for inferring deep relationships when single-gene phylogenies yield uncertain results (Park et al., 2023). For example, multi-gene phylogenies based on transcriptomic and genomic data have proven valuable in resolving deep relationships among divergent clades of archigregarines (Lax et al., 2024). Here, we inferred single-gene phylogenies using SSU rDNA sequences and multi-gene phylogenies from transcriptomic and genomic datasets to advance our understanding of the diversity, phylogenetic interrelationships and classification of marine eugregarine parasites. These data also provide evidence for the establishment of 11 new species.

## 2. Materials and methods

### 2.1. Sample collection

Specimens of marine invertebrates were collected from four different locations in British Columbia: Quadra Island (Hyacinthe Bay), Victoria (Ogden Pint), Bamfield (Grappler Islet), and Vancouver (Stanley Park) (Table 1). Some were collected through SCUBA diving (6-12 m deep) and some were collected from the shore during low-tide. Collected animals were either dissected at the Hakai Institute’s Quadra Island ecological observatory located on Quadra Island or transported to the laboratory to be processed at the University of British Columbia (UBC), Canada. Gregarine cells (i.e., trophozoites) were extracted from either the digestive tract or coelomic spaces of the host using a hand-drawn glass pipette and transferred to a glass slide for examination under a differential interference contrast (DIC) microscope. A Sony Alpha a7R IVA digital camera (ILCE-7RM4A) attached to a Zeiss Axio Scope A1 compound microscope was used at the Quadra marine station, and a Zeiss Axiocam 503 colour camera attached to Zeiss Axioplan 2 microscope or an Axiovert 200 inverted scope was used at UBC. Also, a Sony alpha 7RIII digital camera attached to a Leica DM IL LED inverted microscope was used at both the Quadra marine station and at UBC. After imaging, the cells were washed three times in filtered seawater and stored in PCR tubes containing cell lysis buffer for single-cell transcriptomics. Some gregarines were also isolated for single-cell PCR (Park and Leander, 2024a). Host tissue was stored in 95% ethanol before DNA extraction, PCR and barcoding for host identification.

**Table 1.**
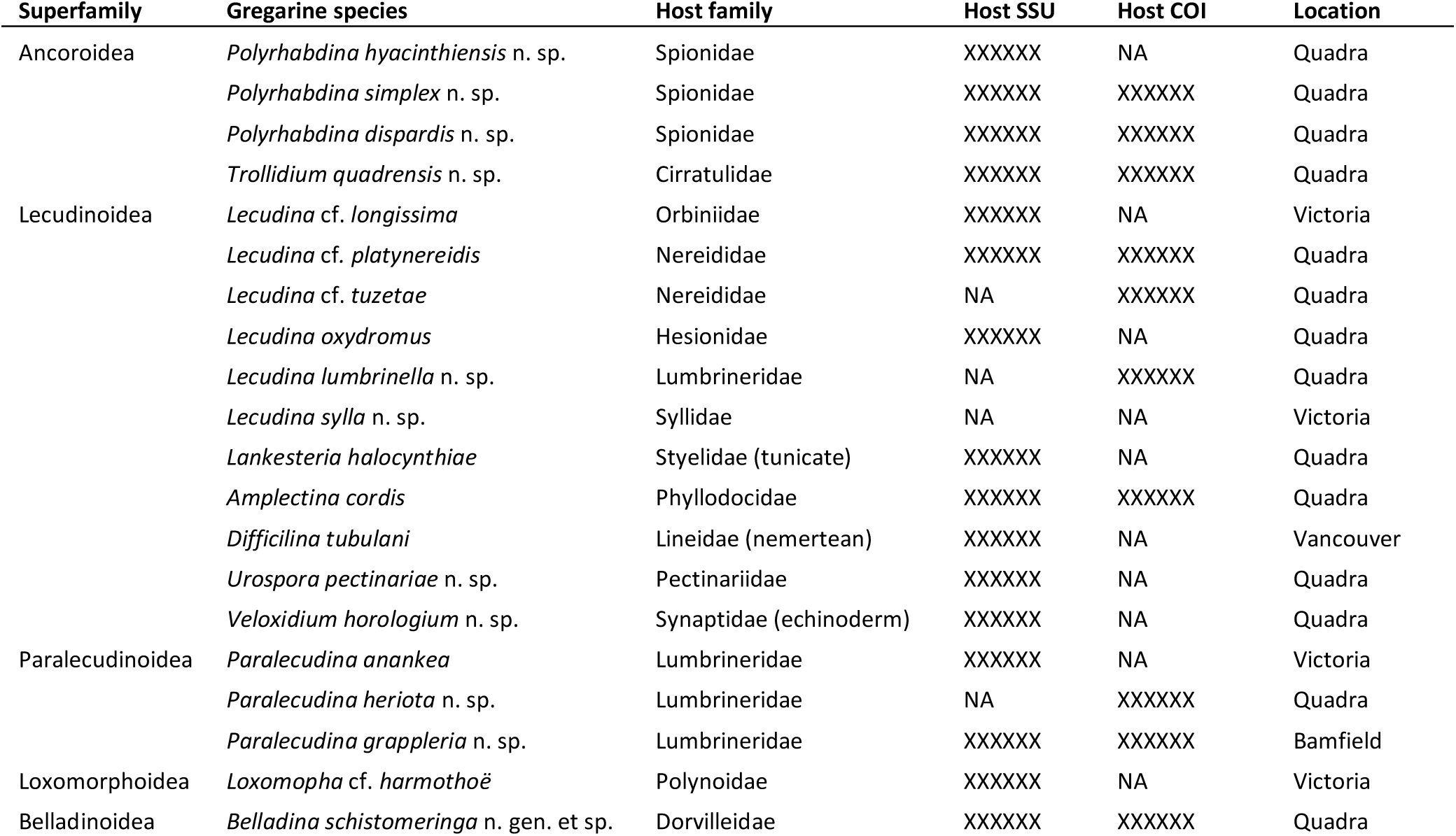
Marine gregarines collected in our study, all from annelid hosts except for three species (noted in parentheses).

### 2.2. Single-cell transcriptomics

Individual trophozoites were kept at −80 °C until they were processed for reverse transcription and cDNA amplification following the Smart-seq2 protocol (Picelli et al., 2014). The cDNA concentrations were measured using a Qubit Fluorometer. Sequencing libraries were constructed using the Illumina Nextera XT protocol and sequenced on an Illumina NextSeq 500 platform with 150 bp paired-end reads by the Sequencing and Bioinformatics Consortium at the University of British Columbia. Errors in the raw reads were corrected with Rcorrector version 1.0.4 (Song and Florea, 2015), and adapters from the Smart-seq2 protocol and Illumina sequencing were removed using Trimmomatic version 0.39 (Bolger et al., 2014). For species with transcriptomes derived from multiple isolates (*Lecudina* cf. *longissima*, *Lecudina sylla* n. sp., and *Loxomorpha* cf. *harmothoe*, *Polyrhabdina dispardis* n. sp.), the trimmed reads were combined prior to assembly. Final transcriptome assemblies were generated using rnaSPAdes version 3.13 (Bushmanova et al., 2019). The completeness of the transcriptomes was evaluated with BUSCO version 5.4.3, using the Alveolata lineage database (Simão et al., 2015) (Supplementary Table 1). Both raw reads and assembled contigs were submitted to the Sequence Read Archive (SRA).

### 2.3. Host identification

Host animals were identified based on their morphology and DNA barcodes by sequencing the partial COI gene and full or near-full-length SSU rRNA gene. Genomic DNA was extracted using the DNeasy Blood and Tissue Kit (Qiagen) following the manufacturer’s protocol. The PCR reaction mixture consisted of 12.3 μl of distilled water, 4 μl of reaction buffer, 0.8 μl of the LCO1490 and HCO2198 primers (Folmer et al., 1994), 0.1 μl of MyTaq polymerase (Bioline), and 2 μl of DNA. PCR was conducted with an initial denaturation at 94 °C for 5 minutes, followed by 35 cycles of 94 °C for 1 minute, 48 °C for 1 minute, and 72 °C for 40 seconds, with a final extension at 72 °C for 10 minutes. PCR products were purified using ExoSAP (Applied Biosystems^TM^) and subsequently sent to the Sequencing and Bioinformatics Consortium at the University of British Columbia, Canada, for sequencing with the same primers used for amplification. In cases where COI sequences could not be obtained, SSU rDNA sequences were retrieved from transcriptome assemblies using barrnap version 0.9 (Seemann, 2013). The host COI and SSU rDNA sequences are deposited in GenBank (see Table 1 for accession IDs).

### 2.4. Phylogenetic analyses of SSU rDNA sequences

The dataset for apicomplexans consisted of SSU rDNA sequences from 135 taxa, including the 20 species with newly obtained transcriptome data in this study, at least one sequence from every known species of marine eugregarine, representatives from other groups of gregarines (i.e., archigregarines, blastogregarines, gregarines from arthropods and terrestrial annelids), representatives from other apicomplexans (e.g., hematozoans, coccidians), and outgroup taxa (e.g., dinoflagellates, chrompodellids, squirmids, ciliates, stramenopiles). Small subunit rDNA sequences were aligned using the E-INS-i algorithm in MAFFT program (Katoh and Standley, 2013) implemented in Geneious Prime. Ambiguous regions were excluded from the SSU rDNA alignments using TrimAl v1.4, resulting in a final alignment of 1,516 bp. Phylogenetic analyses were performed on the CIPRES Science Gateway v3.3 (Miller et al., 2010). Maximum likelihood (ML) phylogenies were constructed using IQ-TREE v2.2.2.7 (Nguyen et al., 2015), with 1,000 replicates for rapid bootstrap analysis, under GTR+F+I+G4 model, which was determined by ModelFinder implemented in IQ-TREE (Kalyaanamoorthy et al., 2017). The best-fitting models of nucleotide evolution for Bayesian analysis was identified using jModelTest v2.1.6 (Darriba et al., 2012), with the corrected Akaike Information Criterion (AICc) selecting GTR+I+G as the optimal model. Bayesian inference was carried out with MrBayes v3.2.7 (Ronquist et al., 2012), running two independent analyses with four chains each for 5 million generations, sampling every 1,000 generations and discarding the first 25% of samples as burn-in. The resulting trees (the best-scoring ML tree and the 50% majority-rule consensus Bayesian tree) were visualized using FigTree v1.4.4 (http://tree.bio.ed.ac.uk/software/figtree/).

### 2.5. Phylogenomic analyses

Open reading frames (ORFs) in the 20 transcriptome assemblies we obtained, along with those from other representative apicomplexans, for a total of 72 (Supplementary Table 2), were identified using TransDecoder v5.5.0 (Haas et al., 2013). The resulting peptide sequences served as input for PhyloFisher v1.2.14, a software that includes a curated database of 240 protein-coding genes and a workflow for phylogenomic analyses (Tice et al., 2021). These sequences were then searched for putative homologs of the 240 genes in the database. The identified putative homologs were added to their respective gene alignments and then processed using PREQUAL v1.02 for the removal of non-homologous characters, BMGE v1.1.2 (Criscuolo and Gribaldo, 2010)and trimAL v1.4 (Capella-Gutiérrez et al., 2009) for trimming, and MAFFT v7.455 (Katoh and Standley, 2013) for alignment generation. Alignment assessment for errors was performed using DIVVIER v.1.01 (Ali et al., 2019). The updated alignments were used for single-gene tree reconstruction with RAxML v8.2.12 (Stamatakis, 2014). Each of the 240 single-gene trees was manually verified using ParaSorter, a graphical user interface tool implemented in PhyloFisher (Tice et al., 2021). Only genes with completeness greater than 50% were selected, producing a supermatrix that includes 44,802 amino acids from 142 genes across 72 species (Supplementary Table 3). A maximum likelihood tree was inferred using IQ-TREE with the LG+C60+F+G4 model (Nguyen et al., 2015). Additionally, another tree was inferred using the LG+C60+F+G4+PMSF model, with the LG+C60+F+G4 tree serving as a guide tree to test the robustness of branch supports.

## 3. RESULTS

### 3.1. Previously described species

Twenty species of marine eugregarines were isolated from annelids, a sea cucumber, a nemertean, and a tunicate, collected from four different locations in British Columbia, Canada (Table 1). The identities of previously described species were confirmed based on trophozoite morphology, host identity, and SSU rDNA sequence similarities (>98.5 %). As a result, nine isolates were identified as previously described species: *Lecudina* cf. *longissimi* (98.5 % identical to FJ832157, Figure 1E), *Lecudina* cf. platynereidis (100% identical to PP819648, Figure 1F), *Lecudina* cf. *tuzetae* (98.8 % identical to JF264868, Figure 1G), *Lecudina oxydromus* (100 % identical to PP819650, Figure 1H), *Lankesteria halocynthiae* (99.6 % identical to KR024694, a micrograph was not obtained), *Amplectina cordis* (100 % identical to PP819651, Figure 1K), *Difficilina tubulani* (99.9% identical to FJ832160, Figure 1L), *Paralecudina anankea* (99.4 % identical to KY678216, Figure 1O), and *Loxomorpha* cf. *harmothoe* (98.5 % identical to MF537616, Figure 1R).

**Figure 1.**
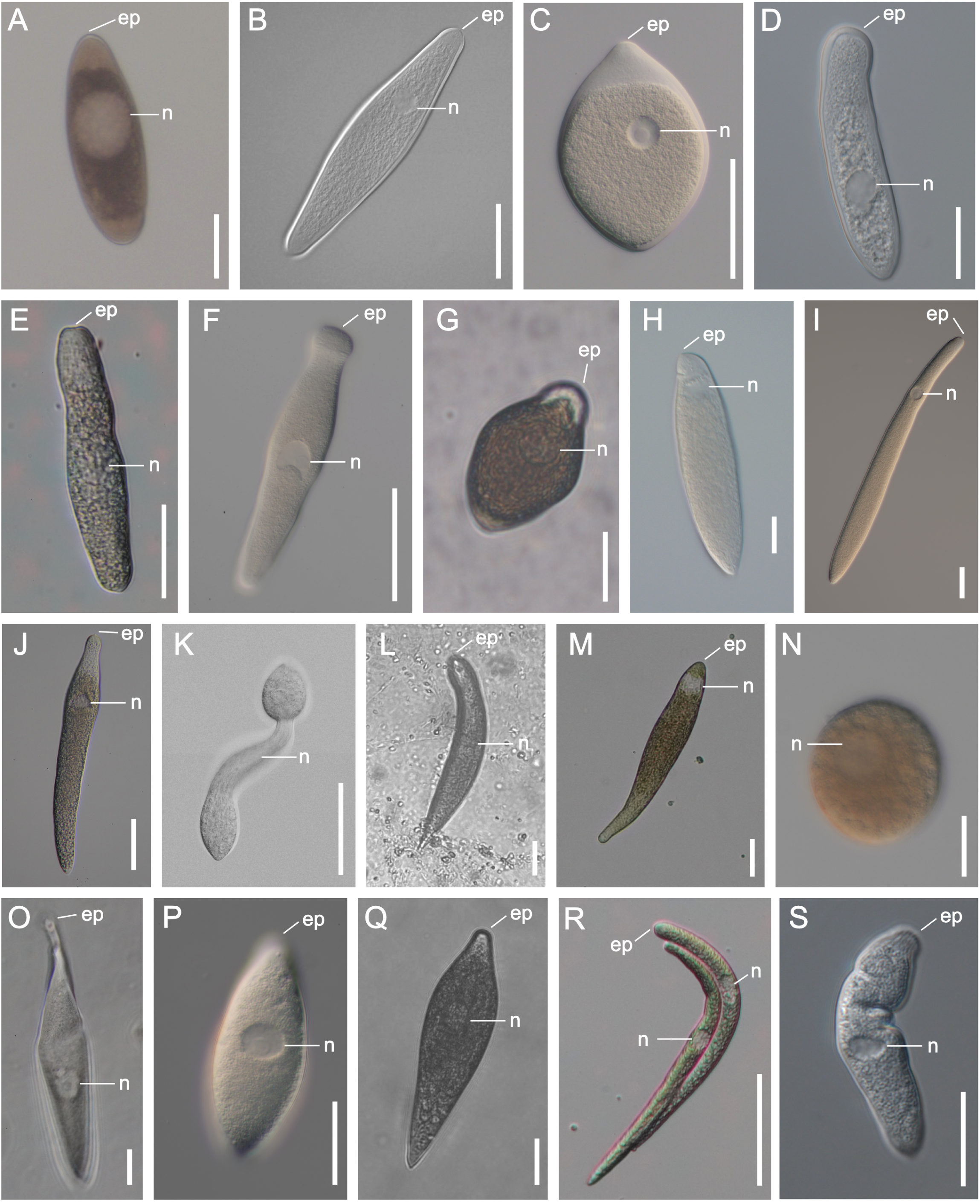
Differential interference contrast light micrographs of marine eugregarine trophozoites from which single-cell transcriptomes were obtained in this study, except for *Lankesteria halocynthiae*. All micrographs show cells with their anterior end oriented upward. **A–D:** The Ancoroidea. (A) *Polyrhabdina hyacinthiensis* n. sp. from a polychaete host (Spionidae gen. sp.), (B) *Polyrhabdina simplex* n. sp. from a polychaete host (Spionidae gen. sp.), (C) *Polyrhabdina dispardis* n. gen. et sp. from a polychaete host (Spionidae gen. sp.), (D) *Trollidium quadrensis* n. sp. from a polychaete host (Cirratulidae gen. sp.). **E–N:** The Lecudinoidea. (E) *Lecudina longissimi* from Orbiniidae gen. sp., (F) *Lecudina* cf. *platynereidis* from a polychaete host (*Platynereis bicanaliculata* species complex), (G) *Lecudina* cf. *tuzetae* from a polychaete host (*Platynereis bicanaliculata* species complex), (H) *Lecudina oxydromus* from a polychaete host (*Oxydromus pugettensis*), (I) *Lecudina lumbrinella* n. sp. from a polychaete host (Lumbrineridae gen. sp.), (J) *Lecudina sylla* n. sp. from a polychaete host (Syllidae gen. sp.), (**K**) *Amplectina cordis* from a polychaete host (Phyllodocidae gen. sp.), (L) *Difficilina tubulani* from a nemertean host (*Lineus torquatus*), (**M**) *Urospora pectinariae* n. sp. from a polychaete host (Pectinariidae gen. sp.), (N) *Veloxidium horologium* n. sp. from a sea cucumber host (Synaptidae gen. sp.). **O–Q:** The Paralecudinoidea. (O) *Paralecudina anankea* from a polychaete host (Lumbrineridae gen. sp.), (P) *Paralecudina heriota* n. sp. from a polychaete host (Lumbrineridae gen. sp.), (Q) *Paralecudina grappleria* n. sp. from a polychaete host (Lumbrineridae gen. sp.). **R:** The Loxomorphoroidea. (R) Two trophozoites of *Loxomorpha* cf. *harmothoe* from a polychaete host (Polynoidae gen. sp.). **S:** The Belladinoidea. (S) *Belladina schistomeringa* n. gen. et sp. from a polychaete host (*Schistomeringos longicornis*). ep, Epimerite; n, nucleus. Scale bars = 40 µm.

### 3.2. Trophozoite morphology of the novel species

#### 3.2.1. Ancoroidea

##### 3.2.1.1. Polyrhabdina hyacinthiensis n. sp. (Figure 1A)

A single spindle-shaped trophozoite was observed, measuring 128 μm long and 47 μm wide (n = 1). Both the anterior and the posterior ends were rounded and devoid of granules. The large, round nucleus measured 36 µm in diameter and was located near the center of the trophozoite. Gliding motility was observed.

##### 3.2.1.2. *Polyrhabdina simplex* n. sp. (Figure 1B)

Trophozoites had an elongated rhombus shape, measuring 113 μm long (n = 12, range = 80-156 μm) and 32 μm wide (n = 12, range = 28-38 μm). A round nucleus, measuring 11 µm in diameter was located between the midpoint and the anterior part of the cell. Both the anterior and the posterior ends were rounded. Gliding motility was observed.

##### 3.2.1.3. *Polyrhabdina dispardis* n. sp. (Figure 1C)

The trophozoites were oblong, slightly curved, 75 μm long and and 49 μm wide (n = 1). The anterior region was devoid of granules and formed a rounded point. The round nucleus was 11 μm in diameter and positioned in the middle of the cell. Gliding motility was observed.

##### 3.2.1.4. *Trollidium quadrensis* n. sp. (Figure 1D)

The trophozoites had an elongated shape, measuring 140 μm long (n = 2, range = 139-141 μm) and 33.5 μm wide (n = 1, range = 32-35 μm). The anterior end was rounded and blunt, while the posterior end was slightly pointed. The round nucleus was 20 µm in diameter (n = 2) and positioned between the midpoint and the posterior end of the cell. Gliding motility was observed.

#### 3.2.2. Lecudinoidea

##### 3.2.2.1. *Lecudina lumbrinella* n. sp. (Figure 1I)

The trophozoites had an elongated shape, measuring 380 µm long and 30 µm wide (n = 1). The posterior part of the cell was wider than the anterior part. The rounded nucleus was located between the anterior end and the midpoint of the cell. The anterior end was rounded and the posterior end was slightly pointed. Gliding motility was observed.

##### 3.2.2.2. *Lecudina sylla* n. sp. (Figure 1J)

The trophozoites had an elongated shape, measuring 130 µm long (n = 4, range = 107-163 µm) and 20 µm wide (n = 4, range = 16-28 µm). The nucleus was 9 µm in diameter and was located near the anterior end at approximately 1/5 to 1/6 of the cell length. The anterior part of the cell was devoid of granules. The anterior end was generally rounded with a papilla (an extruded structure at the tip of the epimerite) at the tip. The posterior end was pointed. Gliding motility was observed.

##### 3.2.2.3. *Urospora pectinariae* n. sp. (Figure 1M)

The trophozoites had an elongated shape, measuring 218 µm long and 33 µm wide. The rounded nucleus was located near the anterior end of the cell. The trophozoites exhibited peristaltic movements, with cytoplasmic contents moving rhythmically along the longitudinal axis, but mostly near the posterior end.

##### 3.2.2.4. *Veloxidium horologium* n. sp. (Figure 1N)

The trophozoite had a spherical shape with a diameter of 86 µm (n = 1); therefore, no anterior or posterior ends were distinguishable. The trophozoite had a round nucleus with a diameter of 32 µm. The cell exhibited peristaltic movement, with cytoplasmic contents moving rhythmically in a clockwise direction.

#### 3.2.3. Paralecudinoidea

##### 3.2.3.1. *Paralecudina heriota* n. sp. (Figure 1P)

Fusiform-shaped trophozoites were 105 µm long and 42 µm wide (n = 1). The ovoid nucleus was located in the center of the cell and was 17 µm long and 21 µm wide. Gliding motility was observed.

##### 3.2.3.2. *Paralecudina grappleria* n. sp. (Figure 1Q)

Fusiform-shaped trophozoites were 218 µm long and 56 µm wide (n = 1). The round nucleus was 18 µm in diameter. Gliding motility was observed.

#### 3.2.4. Belladinoidea

*Belladina schistomeringa* n. gen. et sp. (Figure 1S)

Elongated-shaped trophozoites were 100 µm long and 27 µm wide (n = 1). The ovoid nucleus measured 10 µm long and 16 µm wide. Gliding and bending motility was observed.

### 3.3. Phylogenetic relationships inferred from SSU rDNA sequences

Neither the ML nor Bayesian trees inferred from SSU rDNA sequences recovered gregarines or eugregarines as monophyletic groups (Figure 2). Although the deep internal branches were not well-resolved with this data, several major groups of gregarines were identified in both ML and Bayesian trees. The Lecudinoidea was strongly supported (BS = 100, PP = 1), including *Lecudina*, *Lankesteria*, *Amplectina*, *Sphinctocystis*, *Difficilina*, *Urospora*, *Lithocystis*, *Pterospora*, *Undularius*, and *Veloxidium*. The Ancoroidea was not recovered as a monophyletic group, but instead three different well-supported clades were identified: *Ancora*+*Polyplicarium*, *Polyrhabdina*, and *Trollidium*. Although species of *Trichotokara* were closely grouped with members of the Ancoroidea (i.e., *Trollidium*), this relationship lacked statistical support. *Paralecudina* formed its own lineage with maximal support (BS = 100, PP = 1). *Loxomorpha* and *Cuspisella* were grouped together with strong support (BS = 98, PP = 1). The phylogenetic position of *Belladina schistomeringa* n. sp. was also uncertain, although it was grouped with species of *Selenidium*.

**Figure 2.**
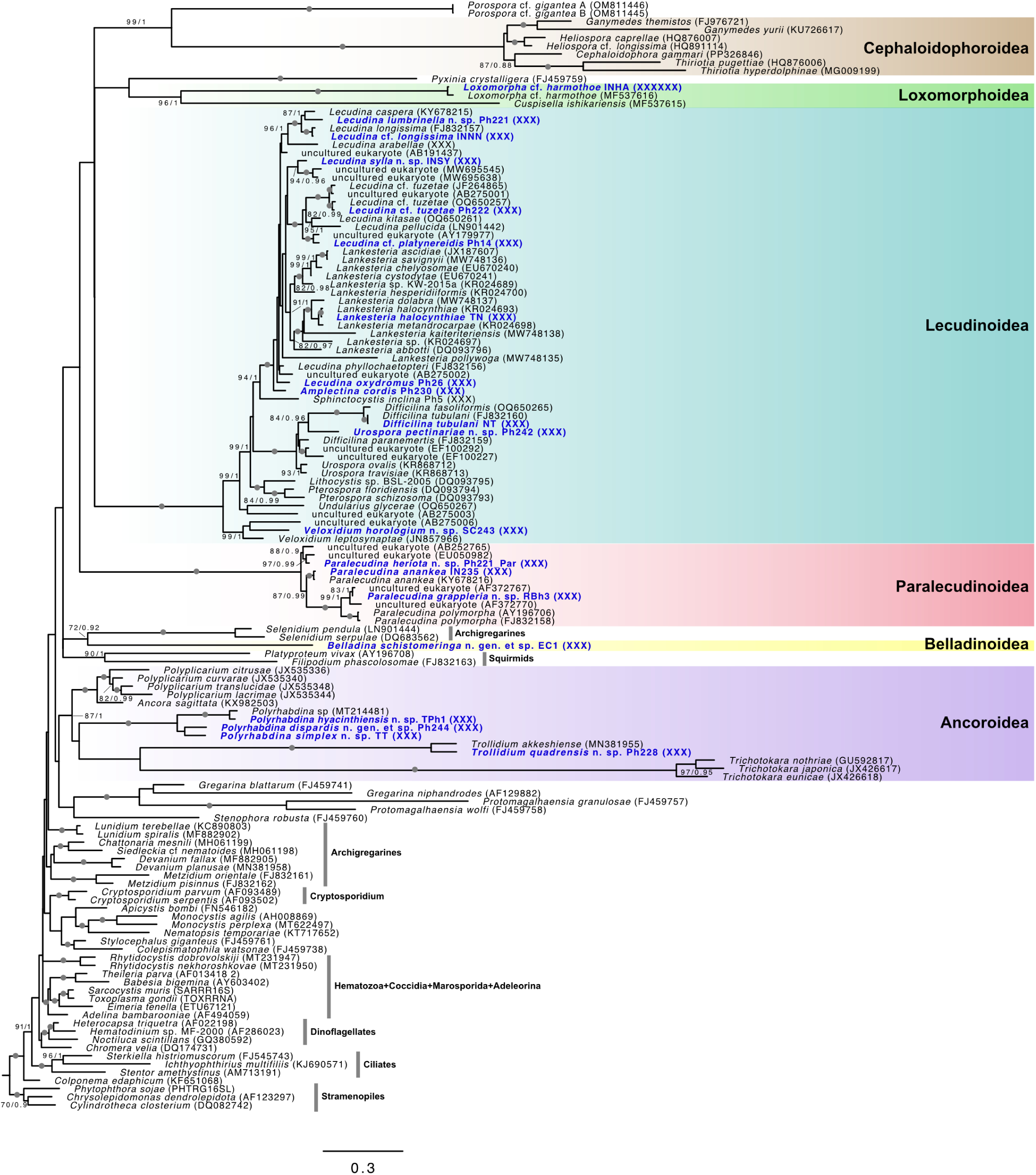
A phylogenetic tree of mostly apicomplexans inferred from 135 SSU rDNA sequences and 1,497 sites using IQ-TREE, generated under the GTR+F+I+G4 model. Bootstrap supports (BS) from IQ-TREE (maximum likelihood) and posterior probabilities (PP) from MrBayes (Bayesian inference) analyses are shown for internal branches. Branches with maximum support in both analyses are marked with grey circles. Taxa with sequences obtained in this study are highlighted in blue font. Major groups of marine eugregarines are shown with colored boxes: The Ancoroidea (purple), Lecudinoidea (blue), Paralecudinoidea (red), Belladinoidea (yellow), Loxomorphoroidea (green), and Cephaloidophoroidea (brown).

### 3.4. Interrelationships of marine eugregarines as inferred from phylogenomic analyses

The Apicomplexa was maximally supported in our ML analyses of the multi-gene dataset (Figure 3). Within the Apicomplexa, the Hematozoa, Coccidia, and Marosporida formed a robust clade (BS = 100), but the position of this clade relative to *Cryptosporidium* and the gregarine clade lacked robust statistical support. All gregarines in the analysis formed a strongly supported monophyletic group (BS = 100) consisting of two major subclades: archigregarines + blastogregarines and eugregarines. Several well-supported subclades were recovered within the eugregarine clade (BS = 100), including the two well-known taxa of marine eugregarines: the Ancoroidea (BS = 100) and Lecudinoidea (BS = 100). Three new superfamily-level clades were also resolved within marine eugregarines: the Paralecudinoidea n. superfam., Loxomorphoidea n. superfam., and Belladinoidea n. superfam. The interrelationships among these marine clades of eugregarines, as well as their relationships to terrestrial clades of eugregarines lacked statistical support.

**Figure 3.**
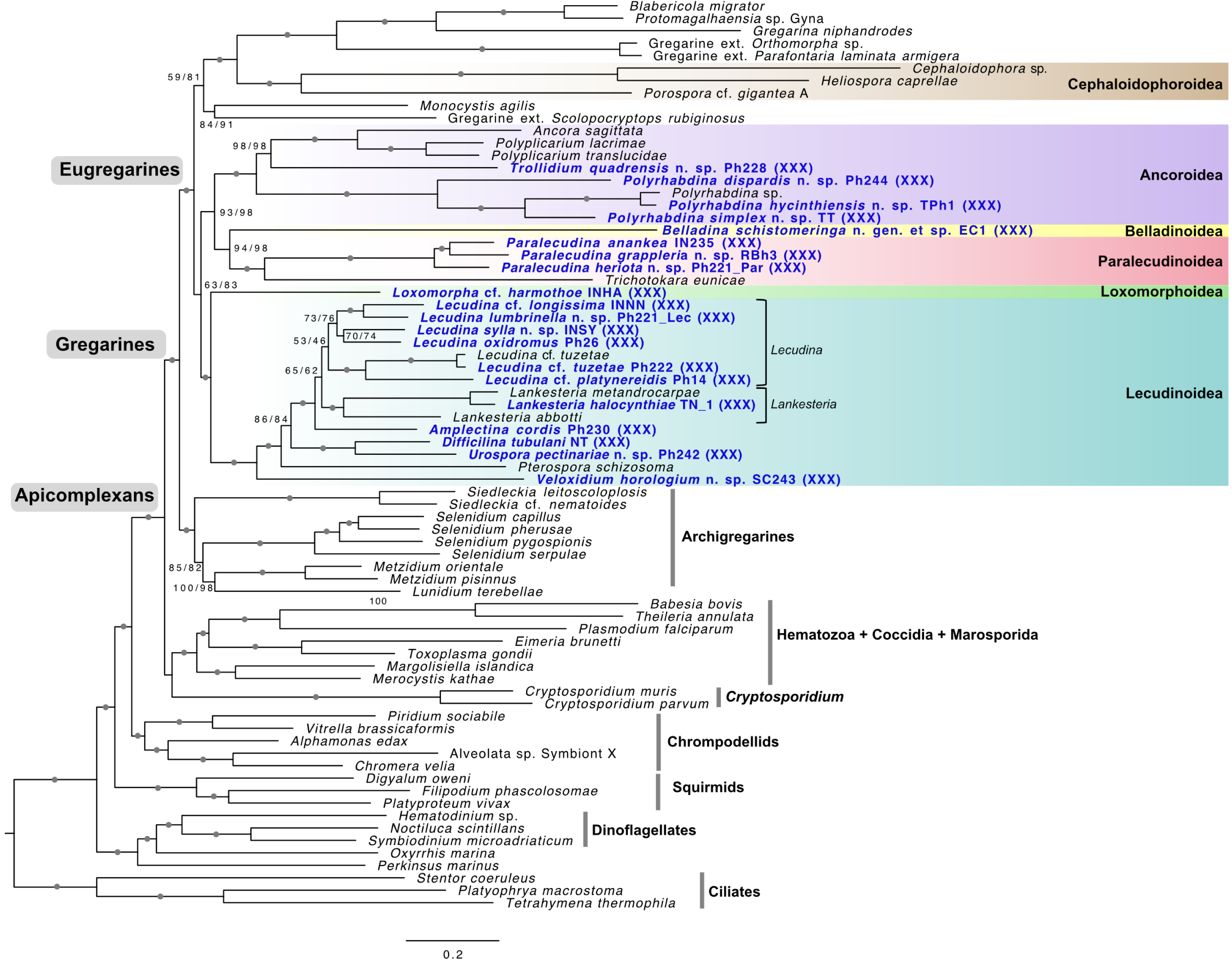
A maximum likelihood phylogenomic tree of alveolates inferred from 44,802 amino acids of 142 genes across 72 species using IQ-TREE, generated under the LG+C60+F+G4 model. For each branch, two bootstrap support (BS) values are shown: one from 1,000 ultrafast bootstrap replicates in IQ-TREE under LG+C60+F+G4, and one under LG+C60+F+G4+PMSF. Branches with maximum support in both analyses are marked with grey circles. Taxa from which single-cell transcriptomes were obtained in this study are highlighted in blue font. Apicomplexans, gregarines, and eugregarines are resolved as monophyletic groups. Major clades of marine eugregarines are shown with colored boxes: The Ancoroidea (purple), Lecudinoidea (blue), Paralecudinoidea (red), Belladinoidea (yellow), Loxomorphoroidea (green), and Cephaloidophoroidea (brown).

## 4. DISCUSSION

### 4.1. An emerging superfamily system for eugregarine classification

Gregarines have been studied since the 1800s due to their diversity, abundance, and relatively large cell size (i.e., the trophozoites) within the gut lumen and other body cavities in invertebrate hosts. However, their classification based solely on morphological characteristics of trophozoites has proven difficult (Grassé, 1953; Kamm, 1922; Levine, 1977a, 1971; Perkins et al., 2000; Rueckert et al., 2011). Molecular phylogenetic analyses have revealed that morphologically similar trophozoites can be highly divergent at the molecular level (Rueckert et al., 2013, 2011), that trophozoites of the same species can exhibit significant morphological variation associated with developmental stages and environmental conditions (Rueckert et al., 2011), and that multiple species of gregarines often co-infect the same host individual (Wakeman, et al., 2014, Lax et al., 2024, Park and Leander, 2024b). As a result, SSU rDNA sequences have become a valuable tool for species identification, despite their limitations in resolving deeper level phylogenetic relationships (Cavalier-Smith, 2014; Rueckert et al., 2011).

With the increasing availability of molecular phylogenetic data, several superfamilies of eugregarines from marine and terrestrial environments have been identified, and currently, eight are formally recognized (Clopton, 2009; Miroliubova et al., 2025; Simdyanov et al., 2017). By integrating existing molecular datasets with newly obtained single-cell transcriptomes from extensive sampling efforts, we propose three additional superfamilies of marine eugregarines, bringing the total to 11 superfamilies of eugregarines: The Actinocephaloidea Léger, 1892; Stylocephaloidea Clopton, 2009; Dactylophoroidea Miroliubova et al., 2025; Stenophoroidea Clopton, 2009; Gregarinoidea Labbé, 1899; Cephaloidophoroidea Rueckert et al., 2011; Ancoroidea Simdyanov et al., 2017; Lecudinoidea Simdyanov, 2013; Paralecudinoidea n. superfam.; Loxomorphoidea n. superfam.; and Belladinoidea n. superfam.

### 4.2. Phylogenomic analyses reveal 5 superfamilies of eugregarines infecting marine annelids

Among the five superfamilies of marine annelid-infecting eugregarines, only the Lecudinoidea includes species that infect other animal hosts (e.g., tunicates, echinoderms, nemerteans). The remaining four superfamilies consists of species that, to our current knowledge, exclusively infect annelid hosts.

#### 4.2.1. The Ancoroidea Léger, 1892

The Ancoroidea comprises four families: The Ancoridae Simdyanov et al., 2017; Polyplicariidae Kamm, 1922; Polyrhabdinidae Cavalier-Smith, 2014; and Trollidiidae Paskerova et al., 2021. Molecular data is currently available for seven species: *Ancora sagittata*, *Polyrhabdina pygospionis*, *Trollidium akkenshiense*, *Poliplicarium citrusae*, *P*. *lacrimae*, *P*. *translucidae*, and *P*. *curvarae* (Paskerova et al., 2021; Simdyanov et al., 2017; Wakeman, 2020; Wakeman and Leander, 2013). We report four novel species belonging to this group: three new *Polyrhabdina* species and one new *Trollidium* species (Figures 1-3, Table 1). *Polyrhabdina hyacinthiensis*, *P*. *simplex*, and *P*. *dispardis* share 96.4%, 87.6%, and 85.28% SSU rDNA sequence similarity, respectively, with *P*. *pygospionis* (MT214481), the only *Polyrhabdina* species from which molecular data was available prior to this study (Paskerova et al., 2021). The SSU rDNA sequences from our newly described *Trollidium* species is 88.4% identical to *T*. *akkenshiense* (MN381955), the only species within the genus from which molecular data was available prior to this study. Although our trees inferred from SSU rDNA sequences did not recover the Ancoroidea as a monophyletic group because of the position of *Trichotokara* (Figure 2), our analyses of phylogenomic data strongly support its monophyly (Figure 3).

#### 4.2.2. The Lecudinoidea Simdyanov and Diakin, 2013

The Lecudinoidea is an exceptionally diverse group at both the molecular and morphological levels, comprising species that infect annelids, nemerteans, tunicates, and echinoderms. Among its many genera, *Lecudina* and *Lankesteria* alone account for more than 80 described species. Despite their diversity, molecular data for members of the Lecudinoidea remains scarce, and previous molecular phylogenetic analyses of SSU rDNA sequences have shown that neither *Lecudina* nor *Lankesteria* are resolved as monophyletic (Iritani et al., 2021; Park and Leander, 2024a). Notably, in our phylogenomic tree, all three species of *Lankesteria* formed a robust clade, which was the sister to a group consisting of all the *Lecudina* species in the analysis, although the monophyly of *Lecudina* was not strongly supported (Figure 3).

We report four novel species and seven previously described species within the Lecudinoidea spanning five genera: seven species of *Lecudina*, one species of *Amplectina*, one species of *Difficilina*, one species of *Urospora*, and one species of *Veloxidium* (Figures 1-3, Table 1). Among the four novel species, *Lecudina lumbrinella* n. sp. was isolated from an unidentified species of Lumbrineridae and has an SSU rDNA sequence that is 92.04% identical to *Lecudina longissima* (FJ832157). *Lecudina sylla* n. sp. was isolated from a Syllidae gen. sp. and has an SSU rDNA sequence that is 92.93% identical to *Lecudina phyllochaetopteri* (FJ832156). *Urospora pectinariae* n. sp. was isolated from ice cream cone worms (Annelida, Pectinariidae) and has trophozoites with an elongate cell shape and peristaltic movement, where cytoplasm flows along the cell’s longitudinal axis. This new species has an SSU rDNA sequence that is 89.92% identical to *Urospora ovalis* from the annelid *Travisia forbesii* (KR868712). Our phylogenomic analyses grouped *Urospora pectinariae* with *Difficilina tubulani* within a robust clade (BS = 100, PP = 1; Figure 2). *Difficilina* was originally established for gregarines infecting nemerteans and the trophozoites of these species share *Lecudina*-like morphological traits (Rueckert et al., 2010). Because peristaltic movement is a common characteristic of *Urospora* (Levine, 1977b), we establish this species as *Urospora pectinariae* n. sp., although the relationships between species of *Difficilina* and *Urospora* will require further investigation with greater taxon sampling.

*Veloxidium horologium* n. sp. was isolated from a sea cucumber host (*Leptosynapta* sp.) and has an SSU rDNA sequence that is 86.4% identical to *Veloxidium leptosynaptae*, which was previously found in the sea cucumber *Leptosynapta clarki* (Wakeman and Leander, 2012). *Veloxidium leptosynaptae* is vermiform and exhibits rapid bending movements, whereas *V. horologium* n. sp. is spherical and exhibits peristaltic movement. Despite these differences in morphology and motility, both species, along with an environmental sequence, form a distinct clade in our tree inferred from SSU rDNA sequences (Figure 2). Gregarines from sea cucumbers have historically been assigned to the genera *Urospora*, *Lithocystis*, and *Cystobia*, though molecular phylogenetic data for these groups from sea cucumber hosts remain unavailable (Desportes and Schrével, 2013). Species of *Cystobia* also exhibit a spherical morphology but typically occur in syzygy pairs, similar to *Pterospora* (Landers and Leander, 2005; Woodcock, 1906).

Some species of *Urospora* from sea cucumbers exhibit several different morphotypes (Cuénot, 1892; Diakin and Paskerova, 2004), but whether or not they represent the same species remains unconfirmed due to a lack of molecular phylogenetic data. Until additional DNA sequences, particularly from *Cystobia* and *Urospora*, become available, we classify *V*. *horologium* n. sp. within *Veloxidium* based on the molecular phylogenetic position and overall SSU rDNA sequence similarity with *V*. *leptosynaptae* (Figure 2).

#### 4.2.3. The Paralecudinoidea n. superfam

To date, two species of *Paralecudina* and three species of *Trichotokara* have been described: *P*. *polymorpha*, *P*. *anankea*, *T*. *nothriae*, *T*. *eunicae*, and *T*. japonica (Iritani et al., 2018c; Leander et al., 2003; Rueckert et al., 2013). We describe two additional novel species of *Paralecudina* in this study: *P*. *heriota* n. sp. was isolated from a Lumbrineridae species at Bamfield, British Columbia, and *P*. *grappleria* n. sp. was isolated from a different Lumbrineridae species at Quadra Island, British Columbia. In addition, our newly sequenced *P*. *anankea* had an SSU rDNA sequence that was 99.42% identical to the previously described *P*. *anankea* from *Lumbrineris inflata* (KY678216). Given the strongly supported sister relationship between *Paralecudina* and *Trichotokara* in our phylogenomic trees (Fig. 3), we propose a new superfamily, the Paralecudinoidea, to accommodate these genera.

#### 4.2.4. The Loxomorphoidea n. superfam

Two gregarine species are known from scale worm hosts (Annelida, Polynoidae) so far: *Loxomorpha harmothoe* and *Cuspisella ishikariensis* (Hoshide, 1988; Iritani et al., 2018a; Simdyanov, 1996). These two species were grouped together with strong support in our phylogenetic analyses, but have long branches in trees inferred from SSU rDNA sequences (Iritani et al., 2018a). The SSU rDNA sequences of *Loxomorpha* cf. *harmothoe* we found were 98.5 % identical to that of *L*. *harmothoe* (MF537616). In the original species description and in this study, both morphotypes were observed with and without an elongated attachment apparatus (Hoshide, 1988; Simdyanov, 1996). Due to its distinctive trophozoite morphology and isolated phylogenetic position in the tree inferred from SSU rDNA sequences, we propose a new superfamily Loxomorphoidea for gregarines infecting scale worms (i.e., *Loxomorpha* and *Cuspisella*).

#### 4.2.5. The Belladinoidea n. superfam

*Belladina schistomeringa* n. gen. et sp. was isolated from the annelid host *Schistomeringos longicornis* (Dorvilleidae). To date, only one gregarine species has been reported from the Dorvilleidae: *Lecudina staurocephali*, which was found in *Staurocephalus rudolphi* in Italy by Mingaziini (1891) and originally described as *Koellikeria* (= *Koellikerella*) *staurocephali* (Levine, 1976). However, the genus *Koellikeria* was not widely accepted by other authors (Grassé, 1953) and was later synonymized with *Lecudina* by Levine (1976). There is very little information available on this species, and the genus name *Koellikeria* was also used by L. Agassiz (1862) for a group of hydrozoans (now renamed *Koellikerina*). Due to the lack of information on *Koellikeria* and the distinctive phylogenetic position of our identified species in trees inferred from both SSU rDNA sequences and phylogenomics (Figures 2 and 3), we propose the new genus *Belladina* for this species and establish a new superfamily-level group, *Belladinoidea*, to accommodate this genus.

### 4.3. Morphological traits of gregarine trophozoites

Morphological features of trophozoites observed under a light microscope provide limited information for higher-level classification, as many *Lecudina*-like eugregarines look similar but reflect deep evolutionary divergences (Park and Leander, 2024a). Conversely, some genera, such as *Ancora*, *Polyrhabdina*, *Trichotokara*, *Loxomorpha*, *Difficilina*, and even *Filipodium* (Squirmida) have historically been placed within the Lecudinidae, despite their substantial morphological differences (Iritani et al., 2018a; Mathur et al., 2019; Paskerova et al., 2021; Rueckert et al., 2010; Rueckert and Leander, 2010; Simdyanov et al., 2017). Nonetheless, some key morphological characteristics related to attachment, movement, and feeding have been identified through ultrastructural studies of trophozoites, especially when comparing archigregarines with eugregarines (Landers and Leander, 2005; Leander, 2007; Schrével et al., 2016; Simdyanov et al., 2017; Simdyanov and Kuvardina, 2007). For instance, the attachment structures differ significantly between the trophozoites of archigregarines and eugregarines. The trophozoites of archigregarines possess an apical complex that facilitates myzocytotic feeding, which has been lost in eugregarines (Schrével et al., 2016).

Different motility patterns of trophozoites also distinguish different groups of gregarines. Archigregarines are typically more flexible, with fewer longitudinal epicytic folds, enabling them to bend, coil, and twist (Leander, 2008; Schrével, 1971). In contrast, eugregarines generally exhibit dense, closely packed longitudinal epicytic folds, which contribute to their structural rigidity and facilitate their characteristic gliding motility (Mackenzie and Walker, 1983; Valigurová et al., 2013). The trophozoites of some eugregarines, such as *Ancora*, *Loxomorpha*, and *Trichotokara*, possess highly developed and elongated attachment apparatuses with hook-like structures (Paskerova et al., 2021; Rueckert et al., 2013). Some members of the Lecudinoidea exhibit dynamic peristaltic movement, such as *Pterospora* and *Veloxidium*, occupy coelomic spaces rather than the intestinal lumen of their host, and lack an attachment apparatus altogether. Therefore, eugregarines exhibit substantial diversity in surface ultrastructure, attachment strategies, and motility patterns that should provide synapomorphies for each major clade recovered in our phylogenomic analyses. Examining these characteristics with high-resolution microscopy is expected to clarify the defining traits of the major subgroups revealed by phylogenomic data and uncover patterns of character evolution across the diversity of eugregarine parasites.

## 5. Taxonomic summary

Phylum Apicomplexa Levine, 1970

Class Gregarinomorpha Grassé, 1953

Order Eugregarinida Léger, 1900

**Superfamily Ancoroidea Simdyanov et al., 2017**

Genus *Polyrhabdina* Mingazzini, 1891

**Polyrhabdina hyacinthiensis n. sp. Park and Leander**

**Diagnosis.** Gamonts ovoid. Both anterior and posterior ends rounded. Aseptate. Nucleus round, located near the center, slightly toward the anterior end. Gliding motility.

**DNA sequence.** SSU rDNA sequence has been deposited in GenBank (accession ID: XXX)

**Type locality.** Hyacinthe Bay, Quadra Island, British Columbia, Canada (50°6’53’’N, 125°13’29’’W)

**Type habitat.** Marine

**Type host.** Spionidae gen. sp. (accession ID for the host SSU rDNA sequence: XXX)

**Location in host.** Intestine

**Iconotype**. Figure 1A

**Zoobank Registration LSID. XXX**

**Etymology**. “hyacinthiensis” means from (-ensis) Hyacinthe Bay.

***Polyrhabdina simplex n. sp.* Park and Leander**

**Diagnosis.** Trophozoites rhomboid. Aseptate. Both anterior and posterior ends rounded. Nucleus round, located between the midpoint and the anterior part of the cell. Gliding motility.

**DNA sequence.** SSU rDNA sequence has been deposited in GenBank (accession ID: XXX)

**Type locality.** Hyacinthe Bay, Quadra Island, British Columbia, Canada (50°6’53’’N, 125°13’29’’W)

**Type habitat.** Marine

**Type host.** *Scolelepis squamata* (Spionidae) (accession ID for the host COI: XXX)

**Location in host.** Intestine

**Iconotype**. Figure 1B

**Zoobank Registration LSID**. XXX

**Etymology**. “Simplex” meaning “simple” in Latin referring to the simple morphology of this species.

***Polyrhabdina dispardis* n. sp. Park and Leander**

**Diagnosis.** Trophozoites overall oval shape but laterally asymmetrical. Aseptate. Anterior area devoid of granules. Anterior end round and posterior end slightly pointy. Nucleus round, located near midpoint, slightly toward anterior part. Gliding motility.

**DNA sequence.** SSU rDNA sequence has been deposited in GenBank (accession ID: XXX)

**Type locality.** Hyacinthe Bay, Quadra Island, British Columbia, Canada (50°6’53’’N, 125°13’29’’W)

**Type habitat.** Marine

**Type host.** *Laonice* sp. (Spionidae) (accession ID for the host COI: XXX)

**Location in host.** Intestine

**Iconotype**. Figure 1C

**Zoobank Registration LSID. XXX**

**Etymology**. “Dispar” means “unequal” in Latin, and “-dis” is used as a suffix. Dispardis refers to the asymmetrical shape of the trophozoites.

*Trollidium* Wakeman 2020

***Trollidium quadrensis n. sp.* Park and Leander**

**Diagnosis.** Trophozoites elongate. Aseptate. Anterior end bluntly rounded and posterior end slightly pointy. Nucleus round, positioned between midpoint and posterior end. Gliding motility.

**DNA sequence.** SSU rDNA sequence has been deposited in GenBank (accession ID: XXX)

**Type locality.** Hyacinthe Bay, Quadra Island, British Columbia, Canada (50°6’53’’N, 125°13’29’’W)

**Type habitat.** Marine

**Type host.** Cirratulidae gen. sp. (accession ID for the host COI: XXX)

**Location in host.** Intestine

**Iconotype**. Figure 1D

**Zoobank Registration LSID**. XXX

**Etymology**. “Quadr” refers to “Quadra”, the island where the specimen was collected. “-ensis” means “from” in Latin. “Quadrensis” means “from Quadra”.

**Superfamily Lecudinoidea Symdyanov et al. 2017**

*Lecudina* Mingazzini, 1891

***Lecudina lumbrinella* n. sp. Park and Leander**

**Diagnosis.** Trophozoites elongate, aseptate. Anterior end rounded, posterior end slightly pointed. Posterior part wider than anterior. Nucleus round, located between anterior and midpoint. Gliding motility.

**DNA sequence.** SSU rDNA sequence has been deposited in GenBank (accession ID: XXX)

**Type locality.** Hyacinthe Bay, Quadra Island, British Columbia, Canada (50°6’53’’N, 125°13’29’’W)

**Type habitat.** Marine

**Type host.** Lumbrineridae gen. sp. (accession ID for the host COI: XXX)

**Location in host.** intestine

**Iconotype**. Figure 1I

**Zoobank Registration LSID. XXX**

**Etymology**. “Lumbrin-“ refers to the family name of the host, which is “Lumbrineridae”, and “-ella” is a suffix."

***Lecudina sylla* n. sp. Park, Na and Leander**

**Diagnosis.** Trophozoites elongate. Aseptate. Anterior part devoid of granules. Anterior end rounded and posterior end pointed. Papilla at anterior end. Gliding motility.

**DNA sequence.** SSU rDNA sequence has been deposited in GenBank (accession ID: XXX)

**Type locality.** Clover Point, Victoria, British Columbia, Canada

**Type habitat.** Marine

**Type host.** Syllidae gen. sp. **Location in host.** intestine **Iconotype**. Figure 1J

**Zoobank Registration LSID**. XXX

**Etymology**. “Syll-” refers to the family name of the host, which is “Syllidae”, and “-a” is a suffix.

*Urospora* Schneider, 1875

***Urospora pectinariae* n. sp. Park and Leander**

**Diagnosis.** Trophozoites elongate. Aseptate. Nucleus round, located near anterior end. Peristaltic movement.

**DNA sequence.** SSU rDNA sequence has been deposited in GenBank (accession ID: XXX)

**Type locality.** Hyacinthe Bay, Quadra Island, British Columbia, Canada (50°6’53’’N, 125°13’29’’W)

**Type habitat.** Marine

**Type host.** Pectinariidae gen. sp. (accession ID for the host SSU rDNA sequence: XXX)

**Location in host.** uncertain

**Iconotype**. Figure 1M

**Zoobank Registration LSID. XXX**

**Etymology**. “Pectinari-“ refers to the genus name of the host, which is “Pectinaria”, and “-ae” is a suffix meaning “associated with”.

*Veloxidium* Wakeman and Leander 2012

***Veloxidium horologium* n. sp. Park and Leander**

**Diagnosis.** Trophozoite spherical. Aseptate. Nucleus round, located near the center. Peristaltic movement.

**DNA sequence.** SSU rDNA sequence has been deposited in GenBank (accession ID: XXX)

**Type locality.** Hyacinthe Bay, Quadra Island, British Columbia, Canada (50°6’53’’N, 125°13’29’’W)

**Type habitat.** Marine

**Type host.** *Leptosynapta* sp. (Synaptidae) (accession ID for the host SSU rDNA sequence: XXX)

**Location in host.** uncertain

**Iconotype**. Figure 1N

**Zoobank Registration LSID. XXX**

**Etymology**. The name ‘Horologium’ derives from the Latin word for ‘clock,’ referring to the peristaltic movement of the spherical cell.

**Paralecudinoidea n. superfam. this study**

**Remarks.** This superfamily includes *Paralecudina* and *Trichotokara*.

*Paralecudina* Rueckert et al. 2012

***Paralecudina heriota* n. sp. Park and Leander**

**Diagnosis.** Trophozoites fusiform. Aseptate. Nucleus ovoid, centrally located. Gliding motility.

**DNA sequence.** SSU rDNA sequence has been deposited in GenBank (accession ID: XXX)

**Type locality.** Hyacinthe Bay, Quadra Island, British Columbia, Canada (50°6’53’’N, 125°13’29’’W)

**Type habitat.** Marine

**Type host.** Lumbrineridae gen. sp. (accession ID for the host COI: XXX)

**Location in host.** Intestine

**Iconotype**. Figure 1P

**Zoobank Registration LSID. XXX**

**Etymology**. “heriot” refers to the “Heriot Island” near the collection site, and “-a” is a suffix.

***Paralecudina grappleria* n. sp. Park and Leander**

**Diagnosis.** Trophozoites fusiform. Aseptate. Nucleus round. Gliding motility.

**DNA sequence.** SSU rDNA sequence has been deposited in GenBank (accession ID: XXX)

**Type locality.** Hyacinthe Bay, Quadra Island, British Columbia, Canada (50°6’53’’N, 125°13’29’’W)

**Type habitat.** Marine

**Type host.** Lumbrineridae gen. sp. (accession ID for the host COI: XXX)

**Location in host.** Intestine

**Iconotype**. Figure 1Q

**Zoobank Registration LSID. XXX**

**Etymology**. xxxxxxx

**Belladinoidea n. superfam. this study**

***Belladina* n. gen. Park and Leander**

**Diagnosis.** Trophozoites elongate. Aseptate. Gliding and bending motility. Parasites of the annelid family Dorvilleidae.

**Type species.** *Belladina schistomeringa* n. sp. Park and Leander

**Etymology**. “Bella-” means “beautiful” in Latin, referring to the beauty of the host animals. “-dina” is used as a suffix to align with “*Lecudina*”, a common genus of marine eugregarines.”

***Belladina schistomeringa* n. gen. et sp. Park and Leander**

**Diagnosis.** Trophozoites elongate. Aseptate. Nucleus ovoid. Gliding and bending motility observed.

**DNA sequence.** SSU rDNA sequence has been deposited in GenBank (accession ID: XXX)

**Type locality.** Hyacinthe Bay, Quadra Island, British Columbia, Canada (50°6’53’’N, 125°13’29’’W)

**Type habitat.** Marine

**Type host.** *Schistomeringos longicornis* (accession ID for the host COI: XXX)

**Location in host.** Intestine

**Iconotype**. Figure 1S

**Zoobank Registration LSID. XXX**

**Etymology**. “Schistomering-“ refers to the host genus name, which is “*Schistomeringos*”, and “-a” is used as a suffix.

**Loxomorphoidea n. superfam. this study**

This superfamily contains *Loxomorpha harmothoe* and *Cuspisella ishikariensis*, both infecting annelid species of Polynoidae (scaleworms).

## DATA ACCESSIBILITY

Transcriptome raw reads are available in the NCBI SRA (BioProject ID: XXXXXX), and assemblies are available at Mendeley Data, V1 (doi: XXXXXX). Host COI and SSU rDNA sequences are available in GenBank with Accession IDs: XXXXX

## ACKNOLWDGEMENTS

This work was funded by grants to BSL from the Tula Foundation’s Hakai Institute and the National Sciences and Engineering Research Council of Canada (NSERC 2019-03986). PJK was supported by the Gordon and Betty Moore Foundation (https://doi.org/10. 37807/GBMF9201). IN was supported by a National Sciences and Engineering Research Council of Canada Scholarship and a University of British Columbia Four-Year Fellowship.

## AUTHUR CONTRIBUTIONS

**Eunji Park:** Conceptualization, Data curation, Formal analysis, Investigation, Methodology, Visualization, Writing – original draft, Writing – review and editing. **Ina Na:** Data curation, Investigation, Writing – review and editing. **Alana Closs:** Resources, Writing – review and editing. **Kyle Hall:** Resources, Writing – review and editing. **Tyrel Froese:** Resources, Writing – review and editing. **Danja Currie-Olsen:** Resources, Writing – review and editing. **Ondine Pontier:** Resources, Writing – review and editing. **Niels Van Steenkiste:** Resources, Writing – review and editing. **Patrick J. Keeling:** Funding acquisition, Resources, Writing – review and editing. **Brian S. Leander:** Supervision, Conceptualization, Funding acquisition, Resources, Writing – review and editing.

